# PPAR-δ rather than PPAR-γ is likely to be the key modulator of central carbon metabolism in human adipocytes

**DOI:** 10.1101/2025.11.08.687092

**Authors:** Leilei Sun, Martin Wabitsch, Jian Yang, Meena Kishore Sakharkar

## Abstract

Adipogenesis involves adipocyte differentiation and synthesis and storage of fats. PPAR-γ is the master regulator of adipogenesis and regulates genes for adipocyte differentiation, lipogenesis, adipocyte survival and adipokine secretion. Central carbon metabolism (CCM) comprises of three key pathways, glycolysis, tricarboxylic acid cycle and pentose phosphate pathway. CCM utilizes carbon sources to provide energy and building blocks for lipogenesis. Although several targets of PPAR-γ have been identified in the CCM pathways, the exact process of how PPAR-γ modulates adipogenesis via CCM remains elusive. In this study, we used real time-qPCR and metabolic arrays for CCM to understand the effect of PPAR activation by PPAR-γ agonist 15d-PGJ2 on CCM and its role in adipogenesis. Our data show that PPAR-δ is likely the target of 15d-PGJ2 and the key modulator of adipogenesis in human SGBS adipocytes at least under current experimental conditions. Further studies are warranted to fully understand the regulation of CCM in adipocytes.

## Introduction

Central carbon metabolism (CCM) refers to a set of fundamental metabolic pathways, which convert carbon-containing molecules like glucose into energy molecules, such as ATP and NADPH, and precursor molecules, such as acetyl-CoA and α-ketoglutarate. It regulates several important cellular signaling pathways including transcriptional regulation and cell stress responses [1, 2]. The metabolic pathways in CCM consist of glycolysis, tricarboxylic acid cycle (TCA) and pentose phosphate pathway (PPP) [3, 4]. Many metabolites are critical precursor molecules required for lipid, amino acid and nucleotide syntheses [5–7]. CCM dysregulation may lead to various types of diseases including metabolic disorders, diabetes, neurodegenerative disorders and cancer [8–12]. For example, aerobic glycolysis is a hallmark of cancer [13], and inhibitors of aerobic glycolysis, such as metformin and lonidamine, have shown promising anticancer activities [14–16].

Human adipose tissue is an integral part of the body and regulates multiple essential physiological functions such as energy storage and regulation, thermal regulation and hormone secretion. It is distributed mainly underneath the skin and around internal organs [17]. Adipose tissue is composed predominantly of mature adipocytes for energy storage, a stromal-vascular fraction containing cells like preadipocytes and endothelial cells, and an extracellular matrix formed by molecules like fibronectin, laminin and collagens [18, 19]. Adipocytes are produced from preadipocytes via adipogenesis, however, preadipocytes can also differentiate into other types of cells such as macrophages and endothelial cells [20, 21]. Dysfunction of adipose tissue contributes to the development of obesity and insulin resistance [22, 23]. CCM has been shown to play a crucial role in adipogenesis and lipogenesis [24–26], for example, acetyl-CoA produced from the TCA cycle and NADPH produced from PPP are important metabolites required for lipogenesis [27, 28]. Exposure of CCM to metabolism-disrupting chemicals (MDCs) may promote obesity [29–31], and monoisononyl-cyclohexane-1,2-dicarboxylic acid ester (MINCH), the primary metabolite of MDC agent diisononyl-cyclohexane-1,2-dicarboxylate (DINCH), can induce human adipocyte differentiation and affect key metabolic pathways [32]. However, the recent metabolomic study by Goerdeler *et al.* showed neither DINCH nor MINCH possess adipogenic potential towards SGBS cells under physiologically relevant concentrations although MINCH induces lipid accumulation [33].

Peroxisome proliferator-activated receptors (PPARs) are a group of ligand-activated nuclear transcription factors that regulate the expression of a wide variety of genes [34]. Upon activation, they bind to the PPAR-responsive regulatory elements (PPRE) and/or PPAR associated conserved motif (PACM) as obligatory heterodimers with retinoid X receptor [35, 36]. Three PPARs (α, β/δ and γ) have been identified and PPAR-γ is most intensively studied. PPAR-γ is primarily involved in the regulation of lipid metabolism, insulin sensitivity and metabolic homeostasis. As a master regulator of adipogenesis, it regulates genes for lipogenesis, adipocyte differentiation, adipocyte survival and adipokine secretion [37]. However, PPAR-α and PPAR-b/d (also known as PPAR-δ) are important regulators of lipolysis and glucose metabolism, controlling fatty acid oxidation and glucose homeostasis [38, 39]. Due to its critical roles in adipogenesis and lipogenesis, CCM is also regulated by PPAR-γ [37, 40–43]. However, little information is available on how CCM is affected by PPAR-γ activation in pre- and differentiated adipocytes. Therefore, in the current study, we decided to address this important question using real time-qPCR array and metabolomics by treating pre- and differentiated human SGBS adipocytes with 15d-PGJ2 (PPAR-γ agonist) and GW9662 (PPAR-γ antagonist).

## Materials and Methods

### Materials

Human preadipocyte cell line SGBS (Simpson–Golabi–Behmel syndrome) was established and validated in coauthor Martin Wabitsch’s laboratory. Cell culture medium, 1:1 mixture of Dulbecco’s modified eagle’s medium with ham’s F12 medium (DMEM/F12, Cat. No. SH30023FS), and Dulbecco’s phosphate-buffered saline (DPBS, Cat. No. SH3002802) were purchased from Hyclone (Cytiva, Vancouver, BC, Canada). Trypsin-EDTA (0.25%, Cat. No. SH30042.02) and Penicillin-Streptomycin (Cat. No. SV30010) were purchased from Hyclone (Logan, UT, USA). Dimethyl sulfoxide (Cat. No. 276855-100 mL), 15d-PGJ2 (1 mg, Cat. No. 538927), biotin (100 mg, Cat. No. B4501), D-pantothenic acid hemicalcium salt (5 g, Cat. No. 21210), apo-transferrin (human, 100 mg, Cat. No. T2252), 3,3′,5- triiodo-L-thyronine sodium salt (T3, 100 mg, Cat. No. T6397), dexamethasone (100 mg, Cat. No. D1756), 3-isobutyl-1-methylxanthine (IBMX, 250 mg, Cat. No. I5879), hydrocortisone (1 g, Cat. No. H0888), bovine serum albumin (BSA, 50 g, Cat. No. A3059), Compete Tablets EASYpack (Roche, Cat. No. 04693116001) and PhosStop EASYpack (Roche, Cat. No. 04906845001) were purchased from Sigma-Aldrich Canada (Oakville, ON, Canada). Rosiglitazone (Cat. No. 71740) was purchased from Cayman Chemical (Ann Arbor, MI, USA). Insulin (Cat. No. 91007C), fetal bovine serum (FBS, cat no. 12483020), bovine calf serum (BCS, Cat. No. 16170), BCA Protein Assay Kit (Cat. No. P123227) and RIPA buffer (Cat. No. TH271222) were purchased from ThermoFisher Scientific (Manassas, VA, USA).

### Culture of SGBS preadipocytes and induction of adipocyte differentiation

The protocol used to culture human SGBS preadipocyte and subsequent induction of differentiation has been previously reported [44]. Briefly, SGBS preadipocyte were cultured in DMEM/F12 containing 10% FBS, 1% penicillin/streptomycin and 1% 5mM biotin/D-pantothenic mixture (biotin: pantothenate= 3.3:1.7) at 37 °C under 5% CO_2_ with cell culture media changed every 2 days. For differentiation, serum free quick-differentiation medium was added to SGBS preadipocytes upon reaching 80-90% confluency. The quick-differentiation medium contains DMEM/F12 supplemented with 1% penicillin/streptomycin, 1% 5 mM biotin/pantothenate mixture, 0.01 mg/mL apo-transferrin, 100 nM cortisol, 0.2 nM T3, 25 nM dexamethasone, 250 µM IBMX, 2 µM rosiglitazone and 20 nM insulin. The quick-differentiation medium was removed on day 4 of differentiation. Then, 3FC medium was added to the cells from day 4 through day 14. The 3FC differentiation cocktail medium consists of DMEM/F12 medium supplemented with 1% penicillin/streptomycin, 1% 5 mM biotin/pantothenate mixture, 0.01 mg/mL apo-transferrin, 20 nM insulin, 100 nM cortisol and 0.2 nM T3. The 3FC medium was not replaced till the cells were harvested.

### Treatments with 15d-PGJ2 and GW9662

Pre- and differentiated human SGBS adipocytes were treated with 100 nM 15d-PGJ2 or 10 μM GW9662 for 24 h before being harvested for subsequent real time-qPCR array and metabolomic studies. Cell culture medium was used as the treatment control.

### Real time-qPCR array

Expression of genes involved in central carbon metabolism was evaluated using the NuRNA Central Metabolism PCR Array (H) from Arraystar Inc. (Rockville, MD, USA) following the manufacturer’s instruction. This array allows for expression profiling of 373 transcripts for the enzymes and proteins in the in the core metabolic pathways and several key transporters. Briefly, RNAs were extracted from treated or control pre- and differentiated SGBS adipocytes and subsequently reverse-transcribed into the first strand cDNAs using the rtStar™ First-Strand Synthesis Kit (Arraystar Inc.). Each cDNA product was diluted in nuclease-free water at 1:80 ratio (mix well and spin down). Then, sample cDNA mix (2010 µL SYBR Green Master Mix, 1600 µL diluted cDNA template and 390 µL ddH_2_O), Genomic DNA Contamination Control (GDC: 5 µL SYBR Green Master Mix, 1 µL no reverse transcriptase control and 4 µL ddH_2_O) and Blank Control Mix (25 µL SYBR Green Master Mix and 25 µL ddH_2_O) were prepared and loaded onto the Central Metabolism PCR array at 10 µL/well. The qPCR was run under the following cycling conditions: initial denaturation (95 °C, 10 min), 40 cycles of cycling (denaturation: 95 °C, 15 sec and annealing, extension, and read fluorescence: 60 °C, 1 min), and melting curve analysis. The obtained amplification data were analyzed using the ΔΔCt method and fold-change for each gene was presented as 2^-ΔΔCt^.

### Metabolomics

The metabolomics was carried out in duplicate at The Metabolomics Innovation Centre (TMIC, Victoria, BC, Canada). Cell pellets (containing ∼10^6^ cells) of pre- and differentiated SGBS adipocytes (control, 15d-PGJ2-treatment and GW9662-treastment) were subjected for Central Carbon Metabolism (93 metabolites) analyses. For each cell pellet, 250-400 µL of water was added and the cells were subsequently lysed on a Retsch MM 400 mill mixer at 30 Hz for 3 min with the aid of two 4-mm metal beads. A 100 µL aliquot of the homogenate from each sample was taken out and transferred to a 1.5 mL Eppendorf tube and 400 µL of methanol-acetonitrile (1:1, v/v) was added. The mixtures were homogenized on the same mill mixer at 30 Hz for 2 min with the aid of two 4 mm metal beads, followed by ultra-sonication in an icy water bath for 3 min and subsequent centrifugal clarification at 21,000 g for 10 min. The clear supernatants were collected for the following analyses by liquid chromatography-multiple reaction monitoring mass spectrometry (LC-MRM/MS). The precipitated pellets were collected and used for protein assay using a standardized Pierce BCA Protein Assay kit according to the manufacturer’s guideline.

### Assay of organic acids

Assay of organic acids of TCA cycle carboxylic acids was carried out using an LC-MRM/MS method that were published previously [45]. Stock solutions containing standard substances of all the organic acids were prepared in methanol at 100 or 500 µM for each compound. The stock solutions were mixed proportionally and then the mixed solution was stepwise diluted to have 10 standard solutions. 20 μL of each standard solution or 20 μL of the clear supernatant of each sample was mixed in turn with 20 μL of an internal standard solution of 10 13C-labeled or deuterated TCA cycle carboxylic acids and 12 deuterated C2 to C22 saturated fatty acids, 80 μL of 200-mM 3-nitrophenylhydrazine solution and 80 μL of 150 mM EDC-6% pyridine methanolic solution. The mixtures were incubated at 50 ℃ for 50 min in a thermomixer at 900 rpm. After reaction, 5 μL aliquots of the resultant solutions were injected into a Waters C18 column (2.1 x 50 mm, 1.7 µm) to run LC-MRM/MS with (-) ion detection on an Agilent 1290 UHPLC system coupled to an Agilent 6495B QQQ mass spectrometer, with the use of 0.01% formic acid in water and 0.01% formic acid in acetonitrile-isopropanol (1:1, v/v) as the binary solvents for gradient elution under optimized conditions.

### Assay of sugars and aldose phosphates

Assay of sugars and aldose phosphates was carried out using an LC-MRM/MS method we published [46]. Briefly, 25 μL of the supernatant of each sample or 25 μL of 10 serially diluted standard solutions of glucose, ribose, ribose-5-phosphate (P), glyceraldehyde -3P, glucose-6P and mannose-6P was mixed with 20 μL of an internal standard solution of 13C-labeled glucose, ribose and glucose-6P, 50 µL of 25-mM 3-amino-9-ethylcarbazole solution, 25 μL of 50 mM sodium cyanoborohydride solution and 10 µL of glacial acetic acid. The mixtures were allowed to react at 60 ℃ for 70 min. After reaction, 150 μL of water and 150 µL of chloroform were added. After vortex-mixing for 15 s and centrifugation for 5 min. 10 µL aliquots of the resultant solutions were injected into a PFP column (2.1 x 150 mm, 1.7 µm) to run UPLC-MRM/MS on an Agilent 1290 UHPLC system coupled to an Agilent 6495B QQQ instrument with positive-ion detection.

### Assay of phosphate-containing metabolites

An internal standard solution containing isotope-labeled ^13^C10-GTP was prepared in 50% methanol. Serially diluted standard solutions of all the targeted metabolites were prepared in the internal standard solution in a concentration range of 0.00001 to 5 µM for each compound. 20 µL of the clear supernatant of each sample was mixed with 80 μL of the internal standard solution. 10 μL aliquots of the sample solutions or the standard solutions were injected into a YMC-Triart C18 column (2.1 x 150 mm, 1.9 µm) to run UPLC-MRM/MS with (-) ion detection on a Waters Acquity UPLC system coupled to a Sciex QTRAP 6500 Plus mass spectrometric instrument, with the use of 5-mM tributylamine solution and acetonitrile as the binary solvents for gradient elution at 0.2 mL/min and 50 °C.

For the above LC-MRM/MS analyses, linearly regressed calibration curves of individual compounds were constructed with internal-standard calibration with the data acquired from standard solutions for each assay. Molar concentrations of the compounds detected in the samples were calculated by interpolating the calibration curves of individual compounds within appropriate concentration ranges, and then normalized with the protein content in mg, which was measured for each sample. Data analyses and comparisons were performed using Metaboanalyst (https://www.metaboanalyst.ca/) online to identify the differentially expressed metabolites (DEM).

## Results

### Changes in central carbon metabolism upon adipocyte differentiation

To evaluate the effects of 15d-PGJ2 and GW9662 on CCM in adipocytes, we first obtained the base level information of CCM in pre- and differentiated human SGBS adipocytes using Real-Time qPCR array and metabolomics. Out of the 373 genes included in the Central Metabolism PCR array, 227 and 10 genes were respectively upregulated and downregulated upon SGBS differentiation using cut-off criteria of |log_2_FC| ≥ 1.00 and *p* < 0.05 (Supplementary Table 1). Out of the 93 metabolites included in the CCM metabolomics, 19 and 50 metabolites were respectively upregulated and downregulated upon adipocyte differentiation using cut-off criteria of |log_2_FC| ≥ 1.00 (Supplementary Table 1). The top 15 upregulated and downregulated genes and metabolites were summarized in Table 1. Genes *PCK1* (log_2_FC = 15.05) and *CECR1* (log_2_FC = -3.53) were the respective most-upregulated and most-downregulated genes, whereas 2-phosphoglyceric acid (log_2_FC = 4.81) and NADH (log_2_FC = -8.62) were the respective most- increased and most-decreased metabolites. Figure 1 (A) shows the volcano plots for differentially expressed genes. Figure 2 (A) and Figure 3 (A) show the Heatmaps for the CCM genes (|log_2_FC| ≥ 1.00 and *p* < 0.05) and metabolites ((|log_2_FC| ≥ 1.00), respectively.

**Figure 1:**
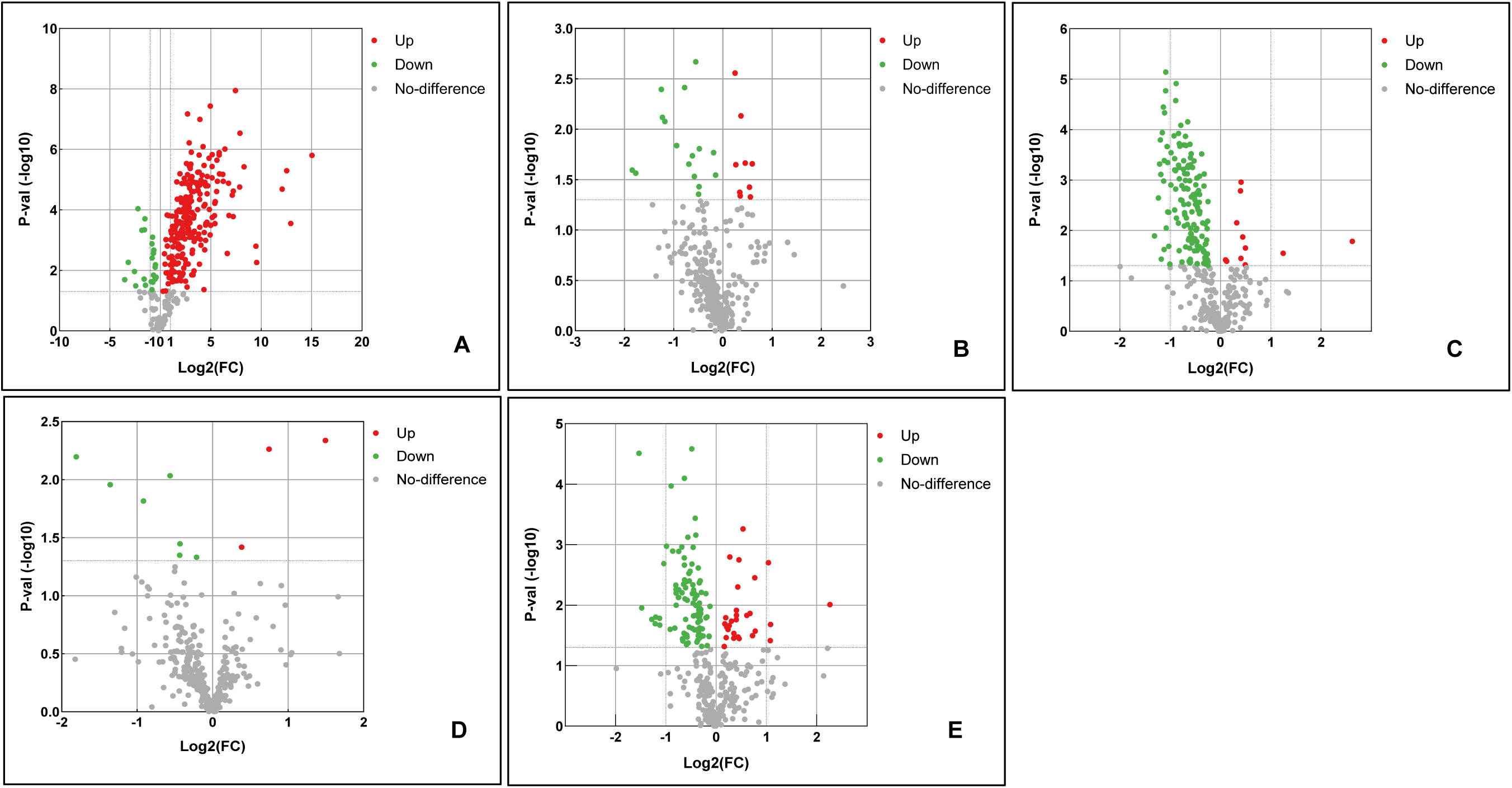
Volcano plots for **A:** Differentiated vs. Pre-adipocytes **B:** Pre-adipocytes treated with 15d-PGJ2 vs. untreated **C:** Differentiated adipocytes treated with 15d- PGJ2 vs. untreated **D:**Pre adipocytes treated with GW9662 vs. untreated **E:** Differentiated adipocytes treated with GW9662 vs. untreated

**Figure 2:**
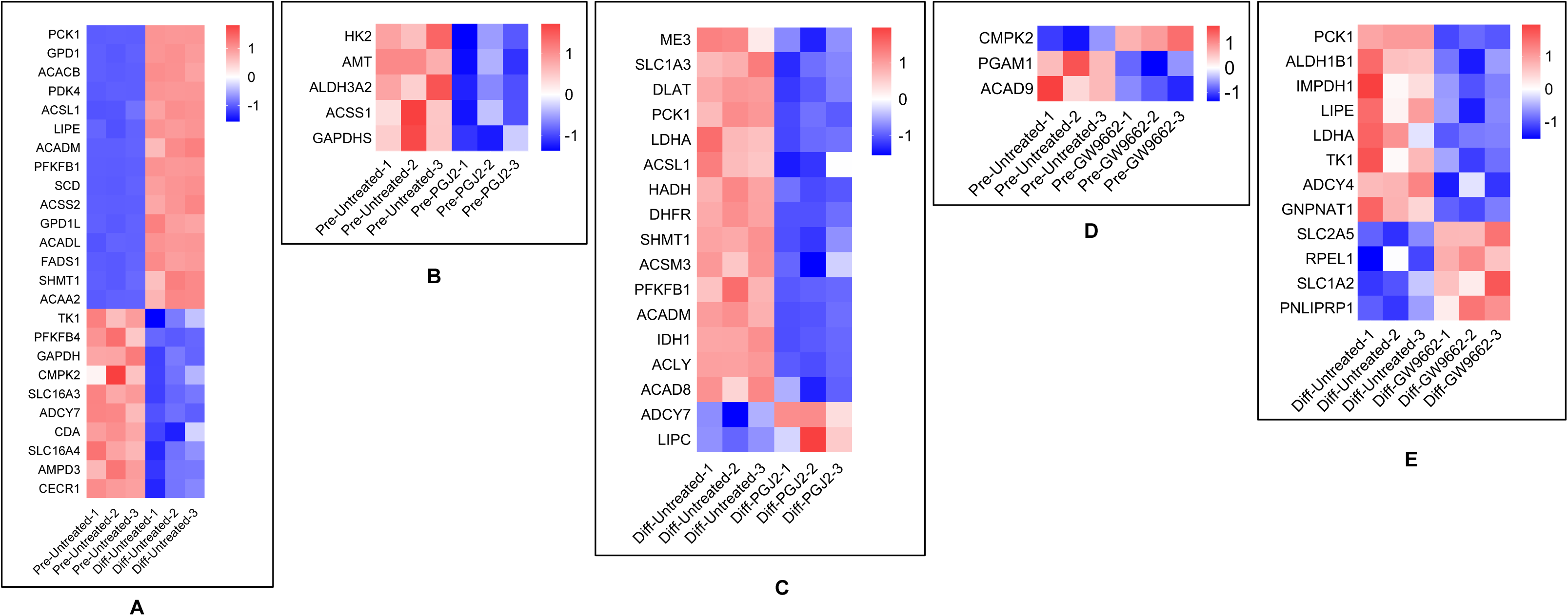
Heatmap for Top 15 upregulated and downregulated CCM genes (|log_2_FC| ≥ 1.00 and p < 0.05): **A:** Differentiated vs. Pre-adipocytes **B:** Pre-adipocytes treated with 15d-PGJ2 vs. untreated **C:** Differentiated adipocytes treated with 15d-PGJ2 vs. untreated **D:**Pre adipocytes treated with GW9662 vs. untreated **E:** Differentiated adipocytes treated with GW9662 vs. untreated

**Figure 3:**
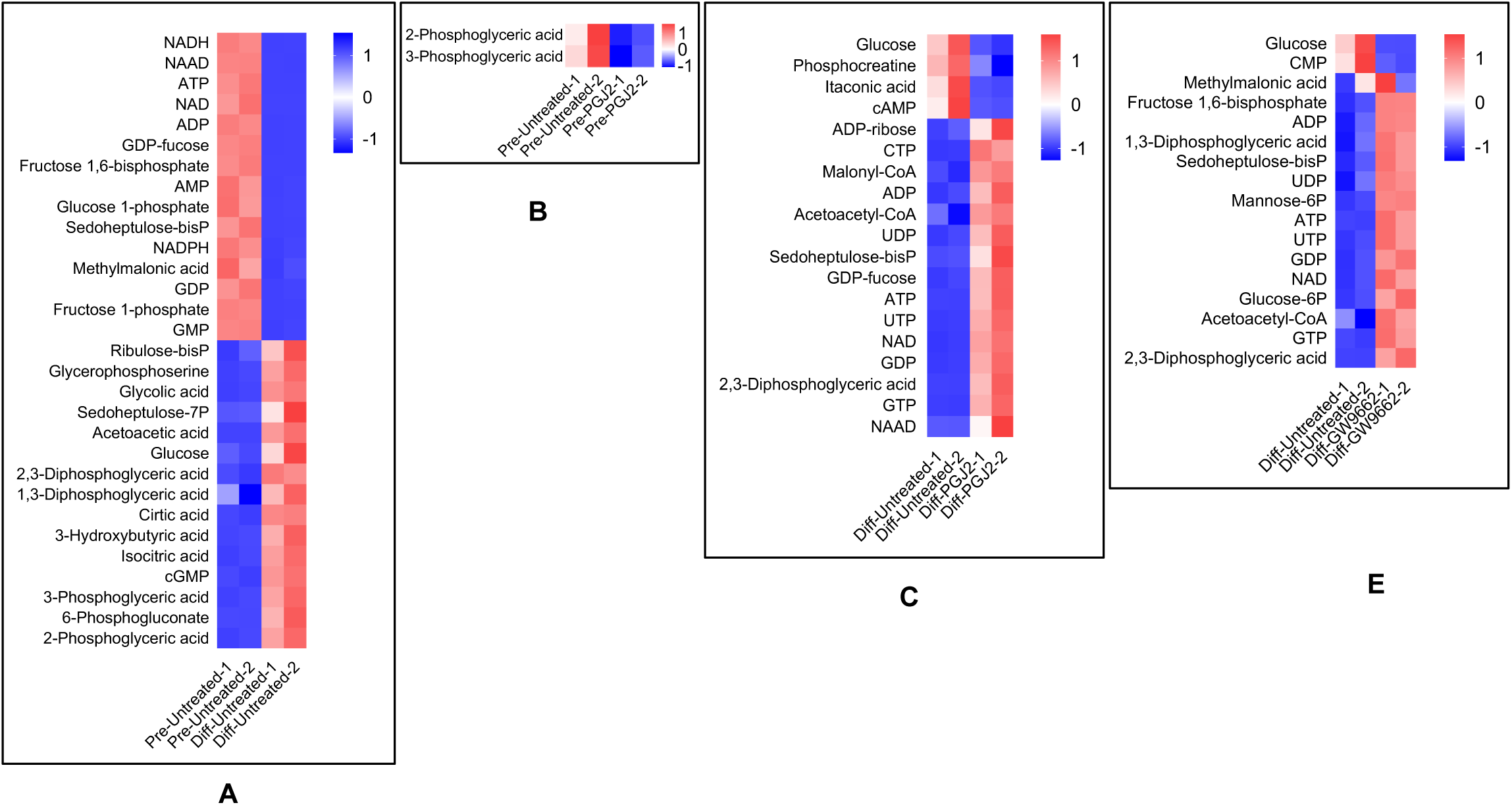
Heatmap for Top 15 upregulated and downregulated CCM metabolites (|log_2_FC| ≥ 1.00): **A:** Differentiated vs. Pre-adipocytes **B:** Pre-adipocytes treated with 15d-PGJ2 vs. untreated **C:** Differentiated adipocytes treated with 15d-PGJ2 vs. untreated **D:**Pre adipocytes treated with GW9662 vs. untreated (No metabolites) **E:** Differentiated adipocytes treated with GW9662 vs. untreated

**Table 1.** Top 15 upregulated and downregulated CCM genes (|log_2_FC| ≥ 1.00 and *p* < 0.05) and metabolites (|log_2_FC| ≥ 1.00) upon SGBS adipocyte differentiation.

| **Upregulated** |  |  |  |
| --- | --- | --- | --- |
| **Genes** | **Log<sub>2</sub>FC** | **Metabolites** | **Log<sub>2</sub>FC** |
| *PCK1* | 15.05 | 2-Phosphoglyceric acid | 4.81 |
| *GPD1* | 12.93 | 6-Phosphogluconate | 4.77 |
| *ACACB* | 12.54 | 3-Phosphoglyceric acid | 4.70 |
| *PDK4* | 12.09 | cGMP | 4.00 |
| *ACSL1* | 9.56 | Isocitric acid | 3.75 |
| *LIPE* | 9.48 | 3-Hydroxybutyric acid | 3.01 |
| *ACADM* | 8.30 | Cirtic acid | 2.73 |
| *PFKFB1* | 7.91 | 1,3-Diphosphoglyceric acid | 2.45 |
| *SCD* | 7.87 | 2,3-Diphosphoglyceric acid | 2.29 |
| *ACSS2* | 7.45 | Glucose | 2.16 |
| *GPD1L* | 7.25 | Acetoacetic acid | 2.06 |
| *ACADL* | 7.23 | Sedoheptulose-7P | 1.98 |
| *FADS1* | 7.14 | Glycolic acid | 1.96 |
| *SHMT1* | 6.81 | Glycerophosphoserine | 1.87 |
| *ACAA2* | 6.76 | Ribulose-bisP | 1.49 |
| **Downregulated** |  |  |  |
| **Genes** | **Log<sub>2</sub>FC** | **Metabolites** | **Log<sub>2</sub>FC** |
| *CECR1* | -3.53 | NADH | -8.62 |
| *AMPD3* | -3.17 | NAAD | -7.02 |
| *SLC16A4* | -2.57 | ATP | -6.63 |
| *CDA* | -2.45 | NAD | -6.55 |
| *ADCY7* | -2.21 | ADP | -6.49 |
| *SLC16A3* | -1.83 | GDP-fucose | -6.09 |
| *CMPK2* | -1.60 | Fructose 1,6-bisphosphate | -5.75 |
| *GAPDH* | -1.57 | AMP | -5.22 |
| *PFKFB4* | -1.53 | Glucose 1-phosphate | -5.14 |
| *TK1* | -1.47 | Sedoheptulose-bisP | -4.91 |
|  |  | NADPH | -4.81 |
|  |  | Methylmalonic acid | -4.72 |
|  |  | GDP | -4.64 |
|  |  | Fructose 1-phosphate | -4.63 |
|  |  | GMP | -4.63 |

### Effects of 15d-PGJ2 and GW9662 on CCM gene expressions in SGBS adipocytes

For SGBS preadipocytes, 15d-PGJ2 treatment downregulated 5 genes *GAPDHS*, *ACSS1*, *ALDH3A2*, *AMT* and *HK2* along with no upregulated genes, whereas GW9662 treatment upregulated gene *CMPK2* and downregulated genes *ACAD9* and *PGAM1* (Table 2). Towards differentiated SGBS adipocytes, 15d-PGJ2 treatment upregulated 2 genes (*LIPC* and *ADCY7*) and downregulated 21 genes (*ME3*, *SLC1A3*, *DLAT*, *PCK1*, *LDHA*, *ACSL1*, *HADH*, *DHFR*, *SHMT1*, *ACSM3*, *PFKFB1*, *ACADM*, *IDH1*, *ACLY*, *ACAD8*, *ME1*, *SDHD*, *PDHX*, *LIPE*, *PGAM4* and *SLC1A6*), whereas GW9662 treatment upregulated 4 genes (*PNLIPRP1*, *SLC1A2*, *RPEL1* and *SLC2A5*) and downregulated 8 genes (*PCK1*, *ALDH1B1*, *IMPDH1*, *LIPE*, *LDHA*, *TK1*, *ADCY4* and *GNPNAT1*) (Table 2). Figure 1 (B-E) shows the volcano plots for differentially expressed genes on treatment with 15d-PGJ2 and GW9662. Figure 2 (B-E) and Figure 3 (B-E) show the Heatmaps for the CCM genes (|log_2_FC| ≥ 1.00 and *p* < 0.05) and metabolites ((|log_2_FC| ≥ 1.00), respectively.

**Table 2.** Differentially expressed CCM genes (|log_2_FC| ≥ 1.00 and *p* < 0.05) upon treatment with 15d- PGJ2 or GW9662 in SGBS pre- and differentiated adipocytes.

| **Preadipocytes** |  |  |  |  |  |  |  |
| --- | --- | --- | --- | --- | --- | --- | --- |
| **Treatment: 15d-PGJ2** |  |  |  | **Treatment: GW9662** |  |  |  |
| Upregulated |  | Downregulated |  | Upregulated |  | Downregulated |  |
| Gene | Log<sub>2</sub>FC | Gene | Log<sub>2</sub>FC | Gene | Log<sub>2</sub>FC | Gene | Log<sub>2</sub>FC |
|  |  | *GAPDHS* | -1.84 | *CMPK2* | 1.49 | *ACAD9* | -1.81 |
|  |  | *ACSS1* | -1.77 |  |  | *PGAM1* | -1.36 |
|  |  | *ALDH3A2* | -1.25 |  |  |  |  |
|  |  | *AMT* | -1.23 |  |  |  |  |
|  |  | *HK2* | -1.18 |  |  |  |  |
| **Differentiated adipocytes** |  |  |  |  |  |  |  |
| **Treatment: 15d-PGJ2** |  |  |  | **Treatment: GW9662** |  |  |  |
| Upregulated |  | Downregulated |  | Upregulated |  | Downregulated |  |
| Gene | Log<sub>2</sub>FC | Gene | Log<sub>2</sub>FC | Gene | Log<sub>2</sub>FC | Gene | Log<sub>2</sub>FC |
| *LIPC* | 2.62 | *ME3* | -1.31 | *PNLIPRP1* | 2.26 | *PCK1* | -1.53 |
| *ADCY7* | 1.25 | *SLC1A3* | -1.23 | *SLC1A2* | 1.08 | *ALDH1B1* | -1.48 |
|  |  | *DLAT* | -1.21 | *RPEL1* | 1.08 | *IMPDH1* | -1.28 |
|  |  | *PCK1* | -1.20 | *SLC2A5* | 1.04 | *LIPE* | -1.21 |
|  |  | *LDHA* | -1.19 |  |  | *LDHA* | -1.20 |
|  |  | *ACSL1* | -1.18 |  |  | *TK1* | -1.12 |
|  |  | *HADH* | -1.16 |  |  | *ADCY4* | -1.12 |
|  |  | *DHFR* | -1.14 |  |  | *GNPNAT1* | -1.04 |
|  |  | *SHMT1* | -1.13 |  |  |  |  |
|  |  | *ACSM3* | -1.12 |  |  |  |  |
|  |  | *PFKFB1* | -1.12 |  |  |  |  |
|  |  | *ACADM* | -1.11 |  |  |  |  |
|  |  | *IDH1* | -1.09 |  |  |  |  |
|  |  | *ACLY* | -1.09 |  |  |  |  |
|  |  | *ACAD8* | -1.08 |  |  |  |  |
|  |  | *ME1* | -1.06 |  |  |  |  |
|  |  | *SDHD* | -1.05 |  |  |  |  |
|  |  | *PDHX* | -1.05 |  |  |  |  |
|  |  | *LIPE* | -1.03 |  |  |  |  |
|  |  | *PGAM4* | -1.03 |  |  |  |  |
|  |  | *SLC1A6* | -1.01 |  |  |  |  |

### Effects of 15d-PGJ2 and GW9662 on levels of CCM metabolites in SGBS adipocytes

Effects of 15d-PGJ2 and GW9662 on CCM in adipocytes were evaluated for 93 metabolites using metabolomics. Towards SGBS preadipocytes, 15d-PGJ2 and GW9662 treatments basically imposed no effects on CCM, with only 2-phosphoglyceric acid and 3-phosphoglyceric acid decreased in concentrations upon 15d-PGJ2 treatment. However, for differentiated SGBS adipocytes, 15d-PGJ2 treatment increased the concentrations of 39 metabolites and decreased the concentrations of 4 metabolites, whereas GW9662 treatment increased the concentrations of 32 metabolites and decreased the concentrations of 2 metabolites (Table 3). NAAD (log_2_FC = 5.69) and glucose (log_2_FC = -3.82) were the most increased and decreased metabolites, respectively, upon 15d-PGJ2 treatment; and 2,3- diphosphoglyceric acid (log_2_FC = 4.63) and glucose (log_2_FC = -2.03) were the most increased and decreased metabolites, respectively, upon GW9662 treatment.

**Table 3.** Differentially expressed CCM metabolites (|log_2_FC| ≥ 1.00) upon treatment with 15d-PGJ2 or GW9662 in SGBS pre- and differentiated adipocytes.

| Preadipocytes: 15d-PGJ2 treatment |  |  |  |
| --- | --- | --- | --- |
| Increased |  | Decreased |  |
| Metabolite | Log<sub>2</sub>FC | Metabolite | Log<sub>2</sub>FC |
| None |  | 2-Phosphoglyceric acid | -1.15 |
|  |  | 3-Phosphoglyceric acid | -1.13 |
| Preadipocytes: GW9662 treatment |  |  |  |
| Increased |  | Decreased |  |
| Metabolite | Log<sub>2</sub>FC | Metabolite | Log<sub>2</sub>FC |
| None |  | None |  |
| Differentiated adipocytes: 15d-PGJ2 treatment |  |  |  |
| Increased |  | Decreased |  |
| Metabolite | Log<sub>2</sub>FC | Metabolite | Log<sub>2</sub>FC |
| NAAD | 5.69 | Glucose | -3.82 |
| GTP | 5.41 | Phosphocreatine | -1.52 |
| 2,3-Diphosphoglyceric acid | 4.90 | Itaconic acid | -1.32 |
| GDP | 4.53 | cAMP | -1.01 |
| NAD | 4.24 |  |  |
| UTP | 4.17 |  |  |
| ATP | 3.78 |  |  |
| GDP-fucose | 3.63 |  |  |
| Sedoheptulose-bisP | 3.63 |  |  |
| UDP | 3.56 |  |  |
| Acetoacetyl-CoA | 2.83 |  |  |
| ADP | 2.74 |  |  |
| Malonyl-CoA | 2.58 |  |  |
| CTP | 2.50 |  |  |
| ADP-ribose | 2.30 |  |  |
| NADH | 2.28 |  |  |
| Hs-CoA | 2.20 |  |  |
| CDP | 2.02 |  |  |
| Fructose 1,6-bisphosphate | 1.98 |  |  |
| 1,3-Diphosphoglyceric acid | 1.96 |  |  |
| Acetyl-CoA | 1.91 |  |  |
| UDP-NAcGal/UDP-NAcGlc | 1.76 |  |  |
| GDP-glucose | 1.55 |  |  |
| GDP-mannose | 1.55 |  |  |
| Glyceric acid | 1.50 |  |  |
| UMP | 1.49 |  |  |
| N-Acetylglucosamine 1-phosphate | 1.45 |  |  |
| Methylmalonic acid | 1.28 |  |  |
| Fructose 1-phosphate | 1.25 |  |  |
| 2-Hydroxyglutaric acid | 1.22 |  |  |
| Glycerol 3-phosphate | 1.18 |  |  |
| Glycerophosphoglycerol | 1.14 |  |  |
| GMP | 1.12 |  |  |
| UDP-glucose/UDP-mannose | 1.10 |  |  |
| cGMP | 1.09 |  |  |
| Pyruvic acid | 1.08 |  |  |
| Ribulose-bisP | 1.07 |  |  |
| Glycerylphosphoethanolamine | 1.04 |  |  |
| AMP | 1.03 |  |  |
| **Differentiated adipocytes: GW9662 treatment** |  |  |  |
| ***Increased*** |  | ***Decreased*** |  |
| Metabolite | Log<sub>2</sub>FC | Metabolite | Log<sub>2</sub>FC |
| 2,3-Diphosphoglyceric acid | 4.63 | Glucose | -2.03 |
| GTP | 3.31 | CMP | -1.05 |
| Acetoacetyl-CoA | 2.72 |  |  |
| Glucose-6P | 2.52 |  |  |
| NAD | 2.40 |  |  |
| GDP | 2.29 |  |  |
| UTP | 2.27 |  |  |
| ATP | 2.08 |  |  |
| Mannose-6P | 2.04 |  |  |
| UDP | 1.78 |  |  |
| Sedoheptulose-bisP | 1.69 |  |  |
| 1,3-Diphosphoglyceric acid | 1.68 |  |  |
| ADP | 1.68 |  |  |
| Fructose 1,6-bisphosphate | 1.54 |  |  |
| Methylmalonic acid | 1.52 |  |  |
| Fructose 6-phosphate | 1.51 |  |  |
| NAAD | 1.50 |  |  |
| GDP-glucose | 1.46 |  |  |
| GDP-mannose | 1.46 |  |  |
| cGMP | 1.42 |  |  |
| CTP | 1.38 |  |  |
| Glyceric acid | 1.36 |  |  |
| UDP-NAcGal/UDP-NAcGlc | 1.24 |  |  |
| 2-Hydroxyglutaric acid | 1.23 |  |  |
| CDP | 1.23 |  |  |
| Glucose 1-phosphate | 1.23 |  |  |
| Ribose | 1.18 |  |  |
| NADH | 1.13 |  |  |
| N-Acetylglucosamine 1-phosphate | 1.11 |  |  |
| Pyruvic acid | 1.09 |  |  |
| Glycerylphosphoethanolamine | 1.05 |  |  |
| Fructose 1-phosphate | 1.05 |  |  |

## Discussion

Adipogenesis includes two phases, determination and terminal differentiation. Determination refers to the commitment of mesenchymal stem cells to become preadipocytes along with the loss of potential for differentiation, whereas terminal differentiation is the process of converting preadipocytes into mature, lipid-filled adipocytes. CCM plays a critical role in adipocyte terminal differentiation by providing building blocks such as acetyl-CoA and malonyl-CoA and energy molecule ATP for subsequent lipogenesis. Thus, understanding the baseline difference is essential to evaluate how PPAR-γ regulates CCM in pre- and differentiated SGBS adipocytes.

As shown in Supplementary Table 1, 227 CCM genes were upregulated, spanning among all three components of CCM, and only 10 CCM genes were downregulated upon SGBS differentiation induced by rosiglitazone. This implicates that CCM was upregulated in differentiated SGBS adipocytes, which is consistent with previous reports [47, 48]. The top 2 upregulated genes were *PCK1* (log_2_FC = 15.05) and *GPD1* (log_2_FC = 12.93), both of which are involved in glycolysis and TCA. Gene *PCK1* encodes phosphoenolpyruvate carboxykinase 1, which catalyzes the conversion of oxaloacetate into phosphoenolpyruvate, CO_2_ and GTP. It is a crucial point of gluconeogenesis, allowing some non-carbohydrate molecules such as proteins and amino acids (specifically aspartate and asparagine) to enter CCM and subsequent lipogenesis. Gene *GPD1* encodes glycerol-3-phosphate dehydrogenase 1, which converts dihydroxyacetone phosphate and NADH into glycerol-3-phosphate and NAD^+^. Glycerol-3- phosphate is a key precursor molecule for lipid synthesis in adipocytes via its conversion into triglycerides [49, 50]. However, upon comparing CCM metabolites between pre- and differentiated SGBS adipocytes, we found that concentrations for most of the metabolites (50 metabolites) decreased after differentiation along with only 19 metabolites elevated their concentrations. Specifically, concentrations of the two major building blocks, acetyl-CoA (log_2_FC = -3.16) and malonyl-CoA (log_2_FC = -4.08), major energy molecule, ATP (log_2_FC = -6.63), and key reducing molecules NADH (log_2_FC = -8.62) and NADPH (log_2_FC = -4.81), significantly decreased. Furthermore, the concentration of NAAD (deamido- NAD, log_2_FC = -7.02), the degraded product of NAADP (nicotinic acid adenine dinucleotide phosphate), also decreased, suggesting elevated glucose uptake in differentiated SGBS adipocytes as NAADP modulates insulin-stimulated and PPAR-γ-induced glucose uptake in adipocytes [51]. All these pieces of evidence implicate that the differentiated SGBS adipocytes were strongly committed to lipogenesis, resulting in decreased levels for most of the CCM metabolites.

All three PPARs are expressed in adipocytes, however, PPAR-γ is the most abundant [52]. PPAR-γ is regarded as the master regulator of adipogenesis and facilitates adipocyte differentiation, lipogenesis, insulin sensitivity and metabolic homeostasis [37, 53], whereas activation of PPAR-α or PPAR-d promotes lipolysis and thermogenesis [54, 55]. CCM plays a critical role in adipogenesis and lipogenesis, however, its regulation by PPAR-γ, especially after adipocyte differentiation, is still far from understood. Thus, in this study, we decided to address this question using real time-qPCR array and metabolomics by respectively treating pre- and differentiated human SGBS adipocytes with 100 nM 15d-PGJ2 (selective agonist PPAR-g, EC_50_ = 2 µM) and 10 µM GW9662 (highly selective PPAR-γ antagonist, IC_50_ = 3.3 nM) [56, 57]. Although 15d-PGJ2 is commonly used at micromolar concentrations during *in vitro* investigations [58, 59], its endogenous intracellular concentrations are significantly lower and only around 1 nM [60]. Thus, we decided to use 100 nM 15d-PGJ2 in the current study, which is likely to be more relevant to physiological conditions. GW9662 is also normally used at low micromolar concentrations in cell studies despite its low IC_50_ towards PPAR-γ [61].

For SGBS preadipocytes, neither 15d-PGJ2 nor GW9662 imposed profound effects on CCM (Tables 2 & 3). Real time-qPCR array showed that 15d-PGJ2 treatment only slightly downregulated five genes (*GAPDHS*, *ACSS1*, *ALDH3A2*, *AMT* and *HK2*) and GW9662 treatment moderately downregulated two genes (*ACAD9* and *PGAM1*) and upregulated one gene (*CMPK2*), whereas metabolomics showed that only two metabolites (2-phosphoglyceric acid and 3-phosphoglyceric acid) were slightly decreased by 15d-PGJ2 treatment and no metabolite was affected by GW9662 treatment. This suggests that CCM might be either insensitive to or not regulated by PPAR-γ in SGBS preadipocytes although it plays a central role in adipogenesis.

For SGBS differentiated adipocytes (Tables 2 & 3), 15d-PGJ2 treatment upregulated 2 genes (*LIPC* and *ADCY7*) and downregulated 21 genes (top three genes: *ME3*, *SLC1A3 and DLAT*), and GW9662 treatment upregulated 4 genes (*PNLIPRP1*, *SLC1A2*, *RPEL1* and *SLC2A5*) and downregulated 8 genes (top three genes: *PCK1*, *ALDH1B1* and *IMPDH1*). Interestingly, no gene was upregulated and genes *PCK1*, *LDHA* and *LIPE* were downregulated by both 15d-PGJ2 and GW9662. No gene was upregulated by 15d-PGJ2 and downregulated by GW9662 or vice versa. However, the metabolomics results showed that 15d-PGJ2 treatment increased the levels of 39 metabolites (top three metabolite:

NAAD, GTP and 2,3-Diphosphoglyceric acid) and decreased the levels of 4 metabolites (top three metabolites: glucose, phosphocreatine and itaconic acid), and GW9662 treatment increased the levels of 32 metabolites (top three metabolite: 2,3-diphosphoglyceric acid, GTP and acetoacetyl-CoA) and decreased the levels of 2 metabolites (glucose and CMP). Concentrations of 27 metabolites (glycerylphosphoethanolamine, pyruvic acid, cGMP, 2-hydroxyglutaric acid, fructose 1-phosphate, methylmalonic acid, N-acetylglucosamine 1-phosphate, glyceric acid, GDP-glucose, GDP-mannose, UDP-NAcGal/UDP-NAcGlc, 1,3-diphosphoglyceric acid, fructose 1,6-bisphosphate, CDP, NADH, CTP, ADP, acetoacetyl-CoA, UDP, sedoheptulose-bisP, ATP, UTP, NAD, GDP, 2,3-diphosphoglyceric acid, GTP and NAAD) were elevated and concentration one metabolite (glucose) was reduced by both 15d- PGJ2 and GW9662. Concentration of no metabolite was decreased by 15d-PGJ2 but increased by GW9662 (with |log_2_FC| ≥ 1.00) or vice versa.

Agonist 15d-PGJ2 and antagonist GW9662 are expected to impose opposite functions on PPAR-γ, however, their treatments elicited similar effects (*i.e.* downregulation of CCM genes and elevation of CCM metabolites) in differentiated SGBS adipocytes. This implies that CCM regulation by 15d-PGJ2 and GW9662 might not be via PPAR-γ but through a different target. Previous studies have identified that 15d-PGJ2 may exert a PPAR-γ-independent function via inhibiting NF-κB [62], and NF-κB function is essential for lipolysis in adipocytes [63]. In addition, BZ-26, a derivative of GW9662, is also capable of inhibiting NF-κB [64]. Thus, 15d-PGJ2 and GW9662 might be able to induce similar responses in differentiated SGBS adipocytes via NF-κB. Since NF-κB plays a critical role in maintaining lipolysis, its inhibition would reduce lipolysis in differentiated SGBS adipocytes. However, the experimental results from both 15d-PGJ2 and GW9662 (downregulate CCM genes and increase CCM metabolites) seem to be opposite from the comparison of CCM between pre- and differentiated SGBS adipocytes (upregulate CCM genes and decrease CCM metabolites), implying that they might stimulate lipolysis and reduce lipogenesis. This makes NF-κB less likely to be a target for their functions. Other than PPAR-γ and NF-κB, another important transcription factor, PPAR-δ, can be affected by 15d-PGJ2 and GW9662. Both

15d-PGJ2 and GW9662 have been identified to be PPAR-δ agonists [65, 66]. PPAR-δ activation triggers lipolysis and suppresses lipogenesis [67, 68], which were consistent with our current observation. Although the less-understood PPAR-α promotes similar functions as PPAR-δ, we have not found any evidence showing that 15d-PGJ2 and GW9662 bind to PPAR-α. Thus, we may conclude that PPAR-δ is likely to be the key modulator of CCM in differentiated adipocytes, at least under current experimental conditions, although PPAR-γ plays a crucial role in adipogenesis and lipogenesis. Our current study provides a new angle in developing strategies to tackle diseases such as obesity and diabetes. Of course, further studies are warranted to fully understand how CCM is regulated in adipocytes.

## Conclusion

PPAR-γ is the master regulator of adipogenesis and lipogenesis. We conducted a multi-omics analysis to understand the role of PPAR-γ in CCM using pre- and differentiated SGBS adipocytes. We observed that differentiated SGBS adipocytes were strongly committed to lipogenesis, resulting in decreased levels for most of the CCM metabolites (shift in flux towards lipid synthesis). Interestingly, on treatment with PPAR-γ agonist 15d-PGJ2 and PPAR-γ antagonist GW9662, SGBS cells showed similar effects - downregulation of CCM genes and increase in the concentration of numerous CCM metabolites. This implies a PPAR-γ independent mechanism for the regulation of CCM by 15d-PGJ2 and GW9662. Since both 15d-PGJ2 and GW9662 can affect PPAR-δ and PPAR-d activation is known to trigger lipolysis and suppress lipogenesis, we suggest that PPAR-δ is the potential target involved in CCM regulation. Further experiments are warranted to understand how CCM is regulated in adipocytes.

